# Enhancing synergy of CAR T cell therapy and oncolytic virus therapy for pancreatic cancer

**DOI:** 10.1101/055988

**Authors:** Rachel Walker, Pedro E. Navas, Samuel H. Friedman, Simona Galliani, Aleksandra Karolak, Fiona Macfarlane, Robert Noble, Jan Poleszczuk, Shonagh Russell, Katarzyna A. Rejniak, Amir Shahmoradi, Frederik Ziebell, Jason Brayer, Daniel Abate-Daga, Heiko Enderling

## Abstract

The poor immunogenicity of pancreatic tumors makes them particularly difficult to treat. Standard chemotherapies and single agent immunotherapies have had notoriously little success in this arena. Oncolytic virus therapy has the potential to enhance the penetration of immunotherapeutically-delivered CAR T cells into the tumor and improve treatment outcomes. We evaluate this potential by combining two different mathematical approaches: an ordinary differential equation model to simulate population level tumor response to cytotoxic activity of T cells, coupled with an agent-based model to simulate the enhancement of CAR T cell penetration by oncolytic virus therapy.

## I. Background

Pancreatic cancer is the fourth most common cause of cancer-related death, with 5–year survival rates of only 7%^[1]^. This poor prognosis is, at least in part, due to the insufficient immunogenicity of pancreatic tumors^[2]^. In recent years, an increasing amount of attention has been given to gene therapy technologies designed to “train” the patient’s immune system to detect and kill cancer cells. This novel approach has the potential to overcome treatment challenges attributable to the strong immunosuppressive measures imposed by both the tumor cell and the stromal microenvironment.

Chimeric antigen receptor (CAR) technology involves the retroviral insertion of genes into primary human lymphocytes, which introduces a highly specific antigen receptor to both direct the cognate anti-tumor interaction and provide the necessary activation signals to elicit a potent tumor-killing response^[3]^. The genetically engineered T cells can be activated *ex-vivo* by antibody-mediated CD3 and CD28 and further expanded massively in culture with cytokines prior to re-infusion into the patient to mount their attack on the cancer cell population. CAR T cell therapy has demonstrated great promise in both the laboratory and the clinical setting^[4]^.

Despite this promise, extravasation of CAR T cells into solid tumors is limited due to the lack of homing cues for *ex vivo* activated T cells, resulting in only an estimated 0.3% of infused cells effectively reaching the tumor^[5]^. Moreover, in pancreatic cancer the presence of fibrotic stroma presents a physical barrier to T cells, impeding their infiltration^[6]^. Improving CAR T cell penetration into the tumor is crucial for increasing treatment efficacy, and may be achievable with complementary local therapy. *In situ* injection of oncolytic viruses has the potential to debulk the tumor and increase stroma permeability for deeper infiltration of CAR T cells at the tumor site, with an additional possibility of further T cell recruitment from the vasculature^[7]^. Attenuated strains of the measles RNA virus have been used for vaccination; oncolytic measles virotherapy has been found to cause substantial death of tumor cells by initial infection and subsequent viral replication within infected cells^[8,9]^.

The augmentation of localized T cell recruitment through oncolytic virus therapy requires the determination of synergy mechanisms. The overall goal of this project is therefore to derive mathematically and confirm experimentally optimal sequencing of oncolytic virus therapy with CAR T cell therapy. Because of the complex dynamics that govern these interactions at different biological scales, this project is necessarily multiscale and multidisciplinary.

## II. Illustative Results of Application of Methods

The first component of the study is the development of a non-spatial, compartmental mathematical model of the dynamics of the respective populations of cancer cells, CAR T cells within the tumor, and CAR T cells circulating in the blood using a system of ordinary differential equations (ODEs). Circulating T cells extravasate into the tumor site where they have the capacity to induce the death of cancer cells until they become exhausted or deactivated.

To represent the influence of oncolytic virus therapy, the cancer cell population was segregated into those infected and those uninfected, with an initial instantaneous transfer of cells into the infected compartment to represent viral injection. When infected, cells undergo death by lysis which can in turn release virus particles from the lysing cell, continuing the spread of the virus through the tumor population. With the addition of oncolytic virus therapy, a reduction in the total tumor cell population was observed; the model predicts synergistic effects of a combination CAR T cell and oncolytic virus therapy, resulting in greater effectiveness than both monotherapies independently (example output demonstrated in Fig. 1).

**Figure 1.**
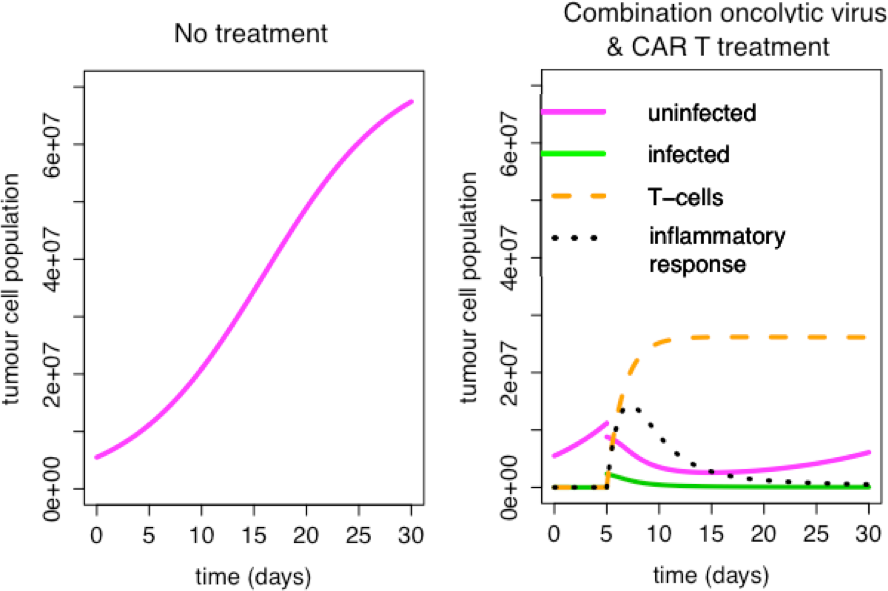
Population dynamics for untreated tumor cells (left) and respective uninfected and infected tumor cells under the influence of synergistic oncolytic virus and CAR T cell therapy (right).

To complement the ODE model, a high-resolution, spatial agent-based model (ABM) was developed that accounts for tumor-CAR T cell interactions and perturbation by oncolytic viruses. We determined the spatial distribution of stroma and cancer cells by discretizing a tissue pathology image using ImageJ^[10]^. The model was able to demonstrate the extravasation of CAR T cells into the tumor site from the defined blood vessels, the initial injection of the oncolytic virus at a predefined tumor site, and the subsequent death of infected cancer cells and corresponding virus spreading. The model allowed simple quantification of the decrease in cancer cell numbers and is configurable for both oncolytic virus and CAR T cell injection timing (an example output can be found in Fig. 2).

## III. Quick Guide to the Methods

### A. Equations

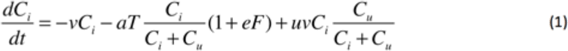

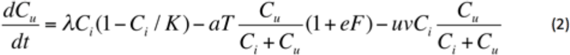

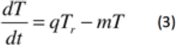

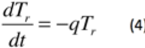

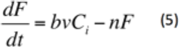

Equations (1) to (5) comprise the ODE model for respective populations of virus-infected cells (*C_i_*), uninfected cells (*C_u_*), T cells in the tumor (*T*) and T cells circulating in the blood (*T_r_*), and the overall inflammatory response (*F*). In (1), the first term on the RHS represents the death of infected cells due to lysis, followed by the T cell killing of cancer cells at a rate proportional to both their interaction rate and the extent of the inflammatory response. The rate of infection of uninfected cells is represented by the last term in (1) and is proportional to the death by lysis. Only the uninfected cancer cells proliferate; growth behavior of the tumor population is obtained from experimental literature and is dependent upon a carrying capacity K. Cells leave the uninfected compartment when they are either killed by T cells or become infected, as previously described. The initial *in situ* virus injection is represented by an instantaneous shift in a subpopulation of the uninfected cancer cells to the infected cell compartment.

**Figure 2.**
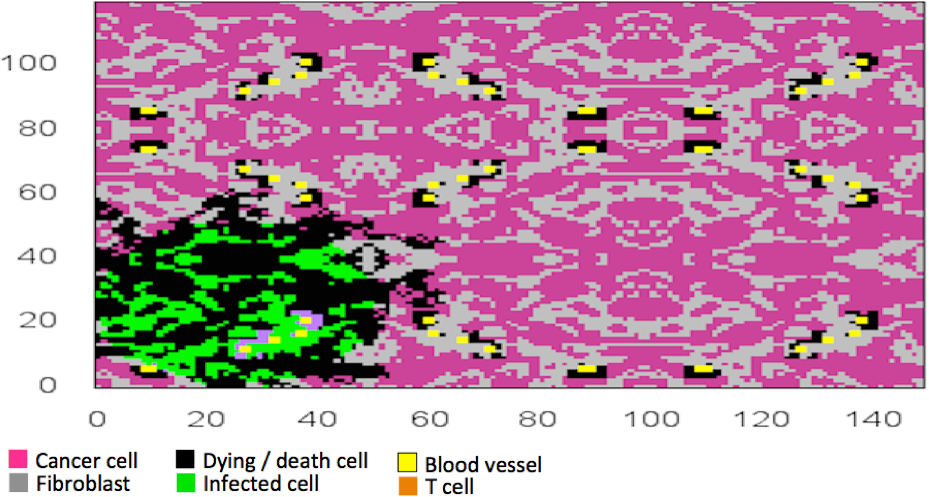
Simulation snapshot of ABM. Note dying cancer cells due to both the oncolytic virus and CAR T cells spreading from the blood vessels.

T cells in the tumor arrive due to extravasation from the blood vessels, and interact with the cancer cells until they become deactivated or exhausted, represented mathematically in (3). The T cell reservoir in the blood is defined by an initial condition representing the instantaneous CAR T cell injection, at which point these activated T cells circulate in the blood and extravasate into the tumor at a fixed rate proportional to their population (4). Finally, in (5), the inflammatory response increases due to death by lysis, and naturally decays.

The model features 8 free parameters, assuming *λ* and *K* the growth rate and carrying capacity of the tumor are derivable from the experimental data. These parameters are the rate of death by lysis (*v*), the rate of T cell killing of cancer cells (*a*), the strength of dependence of T cell killing on the inflammatory response (*e*), the infection rate of uninfected cells (*u*), the infiltration rate of T cells from the blood (*q*), the T cell exhaustion rate (*m*), the dependence of the inflammatory response on death by lysis (*b*), and the rate of decay of the inflammatory response (*n*). While several of these parameters may be obtainable from the experimental literature, further study will allow parameter sensitivity analysis and potential simplification of the model.

Both preliminary models demonstrated the ability to identify conditions for optimizing tumor cell population reduction in combination CAR T cell and oncolytic virus therapies for pancreatic cancer. In future work, we need to more accurately parameterize and calibrate these models to generate robust suggestions for the optimization of treatment protocols. The synergistic outcomes simulated by these models may ultimately be tested in a clinical trial.

## Acknowledgment

We would like to thank Dr. Alexander R. A. Anderson and the Moffitt Cancer Center for organization and support of the 5th Annual Integrated Mathematical Oncology workshop, Immune Cancer, where this project was conceived.

